# A pangenome-graph approach for mapping and imputing barley sequences

**DOI:** 10.64898/2026.08.06.741139

**Authors:** Joan Sarria, Hamza Amhal, Carlos J Ramírez, Ernesto Igartua, Ana M Casas, Bruno Contreras-Moreira

## Abstract

Barley (Hordeum vulgare) is a key cereal crop with exceptional adaptation to diverse environments. With a large, highly repetitive diploid genome, barley presents challenges for pangenome representation. Starting from the reference genome MorexV3, we describe the construction of a barley graph (Pan20) representing the global diversity of landraces and cultivars captured in the public pangenome V1. For mapping arbitrary sequences, a greedy strategy is proposed that combines GMAP alignment followed by intersection with a Practical Haplotype Graph (PHG). This enables presence-absence variation detection and provides a consistent MorexV3 physical coordinate system across genotypes, enabling comparative analysis and visualization. For imputation of genomic data, the PHG approach relies on k-mer pseudo-alignment against the graph. Benchmarks show that Pan20 can accurately align barley genomic and transcriptomic sequences, including those not present in the Morex reference, revealing that a third of long genomic sequences map on non-reference genomes. Moreover, experiments with Genotyping by Sequencing and low-pass sequencing data indicate that FASTQ files can be efficiently mapped and imputed against the graph, preserving local haplotype context. This flexible and scalable graph framework allows barley researchers to explore genetic diversity beyond a single reference and facilitates analysis of diversity panels at the haplotype level, going beyond SNPs. Documentation and a Docker container are available at https://github.com/eead-csic-compbio/barleygraph. The graph sequence mapping utility was added to the Web application https://barleymap.eead.csic.es.

## Introduction

Barley (*Hordeum vulgare* ssp. *vulgare*) remains a foundational model organism and staple crop, occupying a critical role in both global agriculture and genomic research. While domesticated barley and its wild progenitor (*H. vulgare* ssp. *spontaneum*) share a collinear genome, the domestication bottleneck significantly constrained the genetic diversity of cultivated lines (Kilian et al., 2010). A defining morphological divergence between the two subspecies is the acquisition of a tough, non-brittle rachis in domesticated barley, a prerequisite trait that prevented seed dispersal prior to harvest (Pourkheirandish et al., 2015). Concurrently, early domestication events selected for key agronomic mutations, most notably the transition from two-row to six-row spike architecture, the shift toward a spring growth habit (photoperiod insensitivity), and the emergence of naked (hull-less) caryopses (Komatsuda et al., 2007). The accumulation of these adaptive alleles facilitated the geographic expansion of barley across diverse agroecological zones for applications such as human consumption, livestock forage, and malting (Dawson et al., 2015). As the crop disseminated, landraces adapting to localized microclimates became the primary reservoir of genetic diversity for early agrarian societies (Yuan et al., 2025). However, the advent of systematic cross-breeding and mutation induction in the early 20th century catalyzed modern cultivar development, dramatically altering the genetic landscape of the crop. Today, the continued relevance of barley is underscored by its inherent adaptability, the rich allelic repository of its wild ancestor (Wendler et al., 2014), (Milner et al., 2019), and its indispensable contribution to global food security and industrial brewing networks.

The first version of the genome sequence of cultivar Morex paved the way for diverse ways of exploring the diversity of barley germplasm (The International Barley Genome Sequencing Consortium, 2012). This was carried out mostly by calling variants (genomic, exome capture, arrays, etc) in diversity panels with respect to the Morex reference (see for example Bayer *et al*., 2017). An alternative was to use transcriptomic data instead, for instance by clustering transcripts expressed in different genotypes (Contreras-Moreira *et al*., 2017; Rapazote-Flores *et al*., 2019). Examples of Web tools built around the current MorexV3 assembly (Mascher et al., 2021) include GrainGenes, BARLEX, Ensembl Plants, Gramene or BARLEYMAP (Cantalapiedra et al., 2015; Colmsee et al., 2015; Dyer et al., 2025; Olson et al., 2026; Yao et al., 2026). Since the publication of the pangenome V1, comprising 20 landraces and cultivars representing the global diversity pool, the barley community has access to a growing number of annotated genomes that are instrumental to study non allelic variation or gene family expansions (Jayakodi et al., 2020). The Web site https://panbarlex.ipk-gatersleben.de and EoRNA are the primary entry points to these datasets (Milne et al., 2021). However, with these resources come the challenges associated with managing dozens of large genomes that have different physical coordinates, incompatible gene identifiers or divergent annotation (Brůna et al., 2026). Some strategies that have been tested for this include calling pangenes (Contreras-Moreira et al., 2023), building graphs of loci of interest (Jayakodi et al., 2024), graph-based genome browsers (for instance in wheat, Bayer *et al*., 2022) or also pangenome visualizers (http://www.barleypangenome.com, Chapman *et al*., 2026). Despite these advances, modeling a pangenome into a workable graph structure is a non-trivial task; it still requires meticulous optimization of software tools to prevent issues such as unmanageable graph sizes or collapsed repeat regions.

Reference-based algorithms for building pangenome graphs have the advantage that they are easier to update as new genomes become available (Hu et al., 2024). For this reason, in this work we selected a reference-based graph approach that had already been deployed on other crops with complex genomes such as wheat and maize, namely the PHGv2 software (Bradbury et al., 2022). Practical Haplotype Graph (PHG) is able to handle complex genomes through several strategies. First, following the GFF annotation the reference genome is split into genic and intergenic segments and used to define “ranges“; this simplifies the representation of structural variation, as intergenic regions can tolerate large-scale differences without disrupting analyses based on genes. Second, all genomes are aligned to the reference genome (MorexV3 in this case), which provides the whole framework a consistent coordinate system. Unaligned regions spawn new ranges and are assigned MorexV3 positions based on flanking aligned regions. To achieve our goal of building a user-friendly, scalable barley graph we combined a conventional sequence aligner (GMAP) and a precomputed PHG graph (Wu & Watanabe, 2005). This architecture serves two purposes: i) arbitrary sequences can be aligned against the graph genomes, producing global graph coordinates in the MorexV3 space; ii) genotypic data in FASTQ files can be k-mer mapped and imputed in the graph, producing the most likely haplotype string for each sample.

In terms of mapping, our results show that genomic and transcriptomic sequences are effectively aligned and assigned global Pan20 coordinates. While only 2-7% of transcripts match non-reference genomes, about a third of genomic sequences, particularly long reads, align preferably outside MorexV3. We also carried out tests with pangene CDS sequences to measure the accuracy of reported coordinates, finding over 96% exact positions when minimap and GMAP are combined during refinement. We also do a detailed analysis of 28 flowering-related genes to detect and describe the typical alignment challenges in a pangenome, including genes absent in the reference or loci with variable numbers of tandem copies. Finally, our tests with Genotyping by Sequencing (GBS) and low-pass sequencing data indicate that FASTQ files can be efficiently mapped and imputed against Pan20, increasing the fraction of mapped reads and the accuracy of base calls with respect to standard variant calling. In summary, the graph framework built on top of GMAP and PHG proved to be flexible and scalable, allowing barley researchers to explore genetic diversity beyond a single reference.

## Methods

### Building a PHG graph

The Pan20 pangenome graph was built using the PHGv2 framework (release 2.4.74.229) from the data files in **Table S1**. PHG does not construct a full variant graph; instead, it uses a simplified model of genic and intergenic haplotypes (named ranges, defined from MorexV3 anchors) to compactly store genotypes. Genic ranges are padded 500b on each side. Individual genomes are compressed and managed using the AGC version shipping with each PHG release (Deorowicz et al., 2023). A PHG graph is built in several steps (see **Figure S1**): i) the genome assembly (FASTA) and the corresponding gene annotation (GFF) of the reference (MorexV3) are added to define the first set of ranges; ii) additional high-quality assemblies are added one-by-one after alignment to the reference with AnchorWave (Song et al., 2022), adding new ranges; iii) the resulting graph is a Trellis representation mapping ranges across genomes; iv) new assemblies can be added later, requiring only one more AnchorWave alignment per assembly. A critical task carried out in steps i) and ii) is the liftover of reference gene models in each new genome added to the graph. By default in PHG, to establish a consistent coordinate system, all individual assemblies were aligned to the MorexV3 reference using *phg align-assemblies --ref-max-align-cov=1, --query-max-align-cov=1*. Internally this involves extracting CDS sequences with *gff2seq*, which are subsequently lifted over with minimap2 *-x splice -k 12 -a-p 0.4* to define initial anchors (Li, 2018), followed by the *proali* algorithm of AnchorWave. Note that these liftover steps can be performed manually with two aligners (minimap2 [mmap] vs GMAP), and two AnchorWave modes (proali -R 1 -Q 1 [pro] vs genoAli -IV [geno]). In our benchmarks we used minimap2 2.24-r1122, GMAP version 2013-09-30 and AnchorWave v1.2.5. Four graphs were built with the following parameter combinations: mmap_pro (PHG default), mmap_geno, gmap_pro & gmap_geno. Liftover parameters for minimap2 were *-x splice -k 12 -a -p 0.4*; for GMAP *gmapl* they were *-A -f samse*. Graphs are built using *phg create-ref-vcf* and *phg create-maf-vcf,* generating h.vcf and g.vcf for each output .maf from AnchorWave. Merged h.vcf files were generated with *phg merge-hvcfs* and used to compute i) a ML phylogenetic tree with IQTREE v1.6.1 and parameters *-st MORPH -m MK -alrt 1000 -bb 1000* after converting the VCF file to a FASTA file with haplotype alleles encoded as 0-9A-Z characters (Nguyen et al., 2015); and ii) a histogram of range vs genome occupancy with *phgtools core-range-detector*, see https://github.com/jsarriaa/PHGv2Tools. In addition, tables hapIDtable.tsv and hapIDranges.tsv required by *align2graph* (see below) are produced with *phg hapid-sample-table* and *phg sample-hapid-by-range*.

### Barleygraph

The code at https://github.com/eead-csic-compbio/barleygraph was designed around a PHG graph for sequence mapping and haplotype analysis. Inspired by BARLEYMAP (Cantalapiedra et al., 2015), where both genomic and transcript sequences can be fed, sequence alignments are performed with GMAP (Wu & Watanabe, 2005). We used version GMAP v2013-09-30 as it produces smaller indices than more recent versions of GMAP and thus significantly reduces storage requirements. In sequence **mapping mode** (*align2graph,* **Figure S2**), it scans input FASTA sequences hierarchically with GMAP, starting with MorexV3 and moving to the next genome in the list (**Table 1**) until a match satisfying the coverage and identity cut-off values is retrieved. The defaults are 95% and 98%. The coordinates of this alignment are subsequently converted to graph coordinates by overlapping graph ranges with *bedtools intersect* requiring 75% coverage (Quinlan & Hall, 2010). The user can choose to scan any one genome in the graph instead of the default MorexV3 with parameter *--single_genome*. Finally, the coordinates of the corresponding graph ranges are optionally refined (*--add_ranges*) by aligning the input sequences against them. This involves i) cutting the range sequences in the respective genome with AGC, and ii) computing pairwise alignments with GMAP, minimap2 or a combination of both that maximizes coverage. As mentioned in the results, this hybrid strategy might often retrieve positions for sequences not found in the graph but matched by GMAP in individual genomes.

**Table 1.** Number of ranges called by PHG in genomes making up the Pan20 graph depending on the liftover strategy. The last column shows the size of the GMAP index built for each genome.

| genotype | gmap_genom | gmap_pro | mmap_genom | mmap_pro | index (GB) |
| --- | --- | --- | --- | --- | --- |
| MorexV3 | 64,286 | 64,286 | 64,286 | 64,286 | 7.1 |
| Akashinriki | 63,854 | 63,870 | 63,800 | 63,825 | 7.1 |
| B1K-04-12 | 63,706 | 64,002 | 63,626 | 63,936 | 13 |
| Barke | 63,978 | 63,966 | 64,016 | 63,981 | 7.1 |
| Chiba | 63,794 | 63,756 | 63,687 | 63,835 | 7.2 |
| Du_Li_Huang | 63,983 | 64,007 | 63,911 | 64,012 | 7.1 |
| GoldenPromise | 64,046 | 63,949 | 64,097 | 63,955 | 7.1 |
| HOR_10350 | 63,755 | 63,889 | 63,725 | 63,933 | 7 |
| HOR_13821 | 63,944 | 64,020 | 63,866 | 63,981 | 7.1 |
| HOR_13942 | 63,855 | 63,960 | 63,832 | 63,955 | 7.1 |
| HOR_21599 | 64,047 | 64,046 | 63,917 | 63,969 | 7.1 |
| HOR_3081 | 63,883 | 63,641 | 63,894 | 63,624 | 7.1 |
| HOR_3365 | 63,851 | 63,908 | 63,826 | 63,968 | 7.1 |
| HOR_7552 | 63,972 | 63,956 | 63,841 | 63,927 | 7.1 |
| HOR_8148 | 63,887 | 64,052 | 63,789 | 63,959 | 7.1 |
| HOR_9043 | 63,864 | 64,033 | 63,819 | 63,985 | 7.1 |
| Hockett | 63,931 | 64,019 | 63,952 | 64,112 | 7.1 |
| Igri | 63,926 | 64,010 | 63,869 | 64,018 | 7.1 |
| OUN333 | 63,927 | 64,016 | 63,836 | 63,922 | 7.1 |
| Planet | 63,876 | 63,980 | 63,733 | 64,002 | 7.1 |

In **haplotype analysis mode**, sequence reads are mapped against a graph using *phg map-reads* with default parameters except *--min-mem-length*, which should match the length of reads in your FASTQ files. This requires previous indexing with *phg rope-bwt-index* (Li, 2024), which is the most demanding step, taking up to ∼250gb of RAM with Pan20. After mapping, haplotypes are called and imputed with *phg find-paths*, which uses the Viterbi algorithm to solve a hidden Markov model and identify the optimal linear combination of haplotypes from the mapped reads. Diagrams of the resulting haplotypes can be made with *phgtools haplopainting*, producing PNG, PDF or SVG files.

### BARLEYMAP Web application and Docker container

The Web interface of BARLEYMAP was redesigned and upgraded to accommodate the new ‘Graph’ align mode (see **Figure S3** and source code at https://github.com/Cantalapiedra/barleymap_web). In addition to other datasets imported recently, including BaRT2 and panBaRT isoforms, alignments of proteins encoded by barley pangenes from release v14052025 were included from https://github.com/eead-csic-compbio/barley_pangenes (Contreras-Moreira et al., 2023).

The Docker at https://github.com/eead-csic-compbio/barleygraph/pkgs/container/barleygraph can help users analyse haplotypes after downloading precomputed graphs with script *setup_graph*, for instance with parameters *-G Pan20-mmap-pro*. This container can also be used for mapping sequences. In this case GMAP indexes need to be created adding parameter *-g* to the setup script, which will require extra disk space (see **Table 1**).

### Transcriptomic and genomic sequences

Two transcriptomes were used in the mapping benchmark. The *PanBaRT20* dataset compiles transcripts from 20 inbred genotypes representing 19 domesticated and 1 wild barley, the same genomes included in Pan20 (see **Table 1**). It was downloaded from https://ics.hutton.ac.uk/panbart20 and integrates short and long RNA-seq reads from multiple tissues mapped on multiple genome assemblies (Guo et al., 2025). While it annotates isoforms for a total of 79,580 gene models, only the first annotated isoform from each gene was used in the benchmark. The second dataset, named *pantranscriptome16*, mixes long and *de novo* assembled transcripts from cultivars and wild barley (Morex, HarunaNijo, Alexis, AmagiNijo, Beiqing5, Esterel, Franka, Scarlett, SBCC073 Himala2, Hs_ECI-2-0, Hs_Turkey-19-24, Hs_XZ2, Padanggamu, TX9425, Yiwuerleng). It contains 47,588 transcript clusters; only the longest isoform was taken. The FASTA files with the complete transcript sets are available at https://floresta.eead.csic.es/plant-pan-genomes/barley_transcripts and https://github.com/eead-csic-compbio/barleygraph/releases/tag/20260807 (Contreras-Moreira et al., 2017).

In addition, two genomic datasets from cultivar LGDiablo and landrace HOR_495 were included in the mapping benchmark. In the first case, DNA was extracted from a single leaf using the NZY Plant/Fungi gDNA Isolation kit (NZYTech, Portugal) and sent to BMKGENE (Germany), where it was enzymatically fragmented, size-selected (300-600 bp), and prepared using the VAHTS Universal Plus DNA Library Prep Kit (Vazyme, ND801). The library was converted to single-stranded circular DNA using the MGIEasy Universal Library Conversion Kit (MGI) and sequenced on DNBSEQ-T7, producing 2×150bp read pairs. In the second case, the FASTQ file ERR10661231 was downloaded from the European Nucleotide Archive. The HOR495*.HiFi.10K* and *LGDiablo.PE150.10K* FASTA files, available at https://floresta.eead.csic.es/plant-pan-genomes/mapping and https://github.com/eead-csic-compbio/barleygraph/releases/tag/20260807, contain the first ten thousand sequences from the corresponding FASTQ files.

### Barley pangenes

FASTA files of precomputed pangenes were retrieved from https://github.com/eead-csic-compbio/barley_pangenes/releases/tag/19052025, filtering out pangenes containing less than 17 genomes from Pan20 as well as sequences from HOR_10350 due to an assembly version mismatch. The selected CDS nucleotide sequences were mapped with *align2graph --add_ranges [gmap|minimap|both]* and multimapping sequences left out. This experiment was replicated with the two main Pan20 graphs (gmap_geno vs mmap_pro) for several *–minident* values (95, 97 and 100). For gmap_pro and mmap_geno only *–add_ranges both –minident 98* was tested. The analysis consisted of comparing one by one the landing genome start and end coordinates assigned by barleygraph to the known coordinates taken from the pangene FASTA header.

A set of flowering-related pangenes was inspected more carefully to assess the mapping performance across Pan20. These include genes with Presence-Absence (PAV) and Copy Number Variation (CNV), as well as *VRN-H2a* (Horvu_13942_4H01G516500), which is not found in MorexV3 but plays an important role in flowering (Montardit-Tarda et al., 2026). A table with the actual mapped coordinates in all Pan20 genomes is available at https://github.com/eead-csic-compbio/barleygraph/blob/main/paper_Pan20/flowering_bench_Pan20.tsv.

### Imputation tests

Genotyping-by-sequencing (GBS) FASTQ files and genome FASTA files for 25 barley accessions not included in Pan20 (Milner et al., 2019) were obtained from the European Nucleotide Archive (https://www.ebi.ac.uk/ena) and the IPK data archive at https://galaxy-web.ipk-gatersleben.de/libraries/folders/Fd071e794759ab192 (see **Table S5**). Low-pass sequencing files from seven barley cultivars listed in **Table S6** were also tested (Yuan et al., 2025). PHG-based imputation was performed with the Pan20 mmap_pro graph indexed as explained earlier for haplotype analysis, using k-mer mapping (k=31). To this end we ran *phg map-reads --min-mem-length 101* followed by *phg find-paths --path-type haploid*, which generates hVCF files (see **Table S7** and https://phg.maizegenetics.net/hvcf_specifications), and *phg create-fasta-from-hvcf --fasta-type composite*, producing FASTA output.

To evaluate mapping efficiencies, genotyping reads were aligned against three distinct target genomes: i) the MorexV3 reference, ii) the corresponding high-quality (HQ) assembly, and iii) the PHG-imputed composite FASTA file. For i) and ii) alignments were performed using *minimap2 -ax sr -I 8g*. The resulting SAM files were sorted with *samtools sort* and the percentage of successfully mapped reads computed with *samtools flagstat*.

To evaluate error rate, variants resulting from the mappings described above were converted to a MorexV3-based coordinate system. Both the imputed composite and HQ assembly FASTAs were fragmented in 500bp non-overlapping synthetic reads using *bedtools makewindows* and *bedtools getfasta* and subsequently aligned back to the MorexV3 reference using *bwa mem*, followed by *samtools sort*, *bcftools mpileup -T* and *bcftools call -m*. Regarding the MorexV3 GBS variants (i), calls with depth < 10 were filtered out. Artificial heterozygous PHG-imputed calls, introduced while mapping 500bp chunks, were also filtered out. Finally, all three resulting VCF files were joined with *bcftools merge*, genotype fields (GT) extracted with *bcftools query* and compared. Concordance was strictly defined by exact diploid GT equivalence.

## Results

### Building Pan20 pangenome graphs

The Pan20 graph was built as explained in **Figure S1**, starting with MorexV3 and adding the remaining genomes from the barley pangenome V1 iteratively. Overall four versions of the graph were built, depending on the aligner (mmap, gmap) and annotation liftover mode (pro, geno) selected, resulting in different numbers of ranges called in each genome (**Table 1**). **Figure 1** summarizes the Pan20 graph built with minimap2 and the proali mode (mmap_pro), which is the default in PHG. The circular grey plot shows the number of different haplotypes found within ranges, with the lowest values observed in pericentromeric regions. The occupancy barplot inside shows how ranges are found in a variable number of genomes in the graph, with 16,441 found in all (core), 47,672 found in 2 to 19 genomes (accessory) and 173 unique. Finally, the phylogenetic tree at the bottom, rooted on wild genotype B1K-04-12, shows how the different genomes in the graph relate to each other in terms of shared range haplotypes.

**Figure 1.**
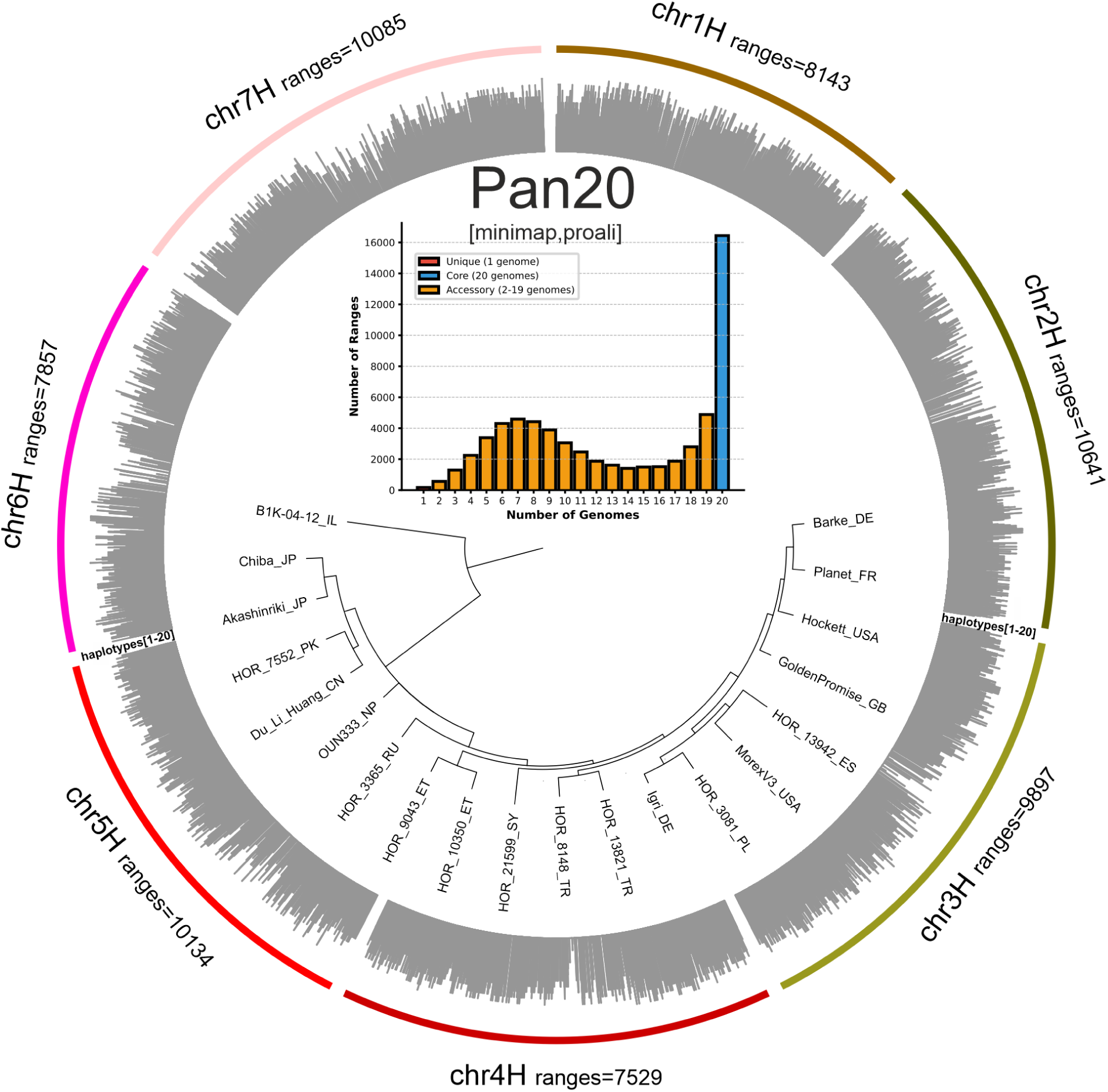
Summary of the Pan20 graph [minimap2, proali]. Outside circle) barplot of the number of distinct haplotypes called by PHG within ranges and chromosomes, with values from 1 to 20. Inside barplot) graph ranges vs number of genomes (pangenome occupancy). Inside tree) Phylogenetic tree of haplotypes found in 64,286 graph ranges, with approximate Bayes branch support values of 100% in all cases except OUN333 (0%) and the JP,PK,CN clade (82%). The country of origin of barley genotypes is added in two-letter code. The dendrogram was plotted with https://tree.bio.ed.ac.uk/software/figtree.

In computing terms, it took a week to build in an ordinary Linux server with 256GB RAM. The bare bones graph is about 4GB large, but the disk requirements grow significantly when the GMAP (150GB) or imputation indices (120GB free disk space, 13GB final files, >12GB RAM) are added.

### Barleygraph leverages alignments and graphs

We developed the Barleygraph utilities as a way of combining the accuracy and speed of alignment tools and the breadth of precomputed pangenome graphs, that efficiently store individual genomic segments included in ranges. In particular, we were concerned with two problems that barley breeders face: the mapping of sequences of interest in a graph with consistent coordinates across genotypes, and the imputation and inference of haplotypes at genome-scale. Accordingly, we developed the sequence mapping and haplotype analysis modes.

The *align2graph* utility was developed for the sequence mapping mode and its main idea is to scan a sequence against the list of genomes making up a graph and take the first reliable match as an entry point to query the graph, retrieve the overlapping range and finally crop and align the individual genome ranges. As shown in more detail in **Figure S2**, the list of genomes is processed, starting with MorexV3, until a good match is found. This means that not all 20 genomes need to be scanned; in fact in most cases only 1-2 are actually searched for hits. The advantage of using a graph is that all sequence matches overlapping ranges can be assigned global graph coordinates, even with sequences unaligned in MorexV3. This utility was added to the BARLEYMAP server at https://barleymap.eead.csic.es (see **Figures S3** and **S4**). The ‘Graph’ button in the Web interface allows users to submit and map genomic and transcriptomic sequences, yielding global graph coordinates and also coordinates of each matching genome. In addition to decorating the output with neighbour markers and genes (such as panBaRT isoforms), flanking MorexV3 genes can be clicked on to inspect multiple alignments of protein sequences of the corresponding pangenes containing them. Users mapping large numbers of sequences are encouraged to use the Barleygraph Docker container instead.

As we expect that the haplotype analysis mode will require reading potentially large FASTQ files we only ship this utility as a Docker container. This way users can analyze their own input files on their local computers.

### Mapping genomic fragments and transcripts

In this section we report on experiments where transcripts and genomic sequence datasets were mapped to Pan20. Only mmap_pro and gmap_geno results are shown, as these were identified as the two competing settings. The goal was to measure how much sequence mapability pangenome graphs provide beyond the reference MorexV3.

**Figure 2A** summarizes the results for transcript mapping. Regarding the *PanBaRT20* dataset, 99.9% of transcripts could be mapped to the Pan20 graph, with most being assigned global graph coordinates (97.9% and 98.4% for mmap_pro and gmap_geno, respectively). Sequences in the *pantranscriptome16* dataset have a lower mapping rate (95.5%), with 91.2% and 90.1% being assigned graph coordinates. Separate mapping tests relaxing the coverage cutoffs revealed a fraction of transcripts actually overlap 2+ ranges and therefore are not assigned graph coordinates with default parameters. Finally, we observed that up to 2% of *PanBaRT20* and 7% *pantranscriptome16* sequences match genomes other than MorexV3. These would have been missed if only the reference genome was used for mapping. One example is the 191449_TR26628-c0_g1_i1 isoform from *pantranscriptome16* (**Table S2**), likely encoded by the *VRN-H2* vernalization locus, that matches the HOR_9043 genome instead of MorexV3, but still is assigned global coordinates chr4H:604188191-604202141 in both mmap_pro and gmap_geno graph variants.

**Figure 2.**
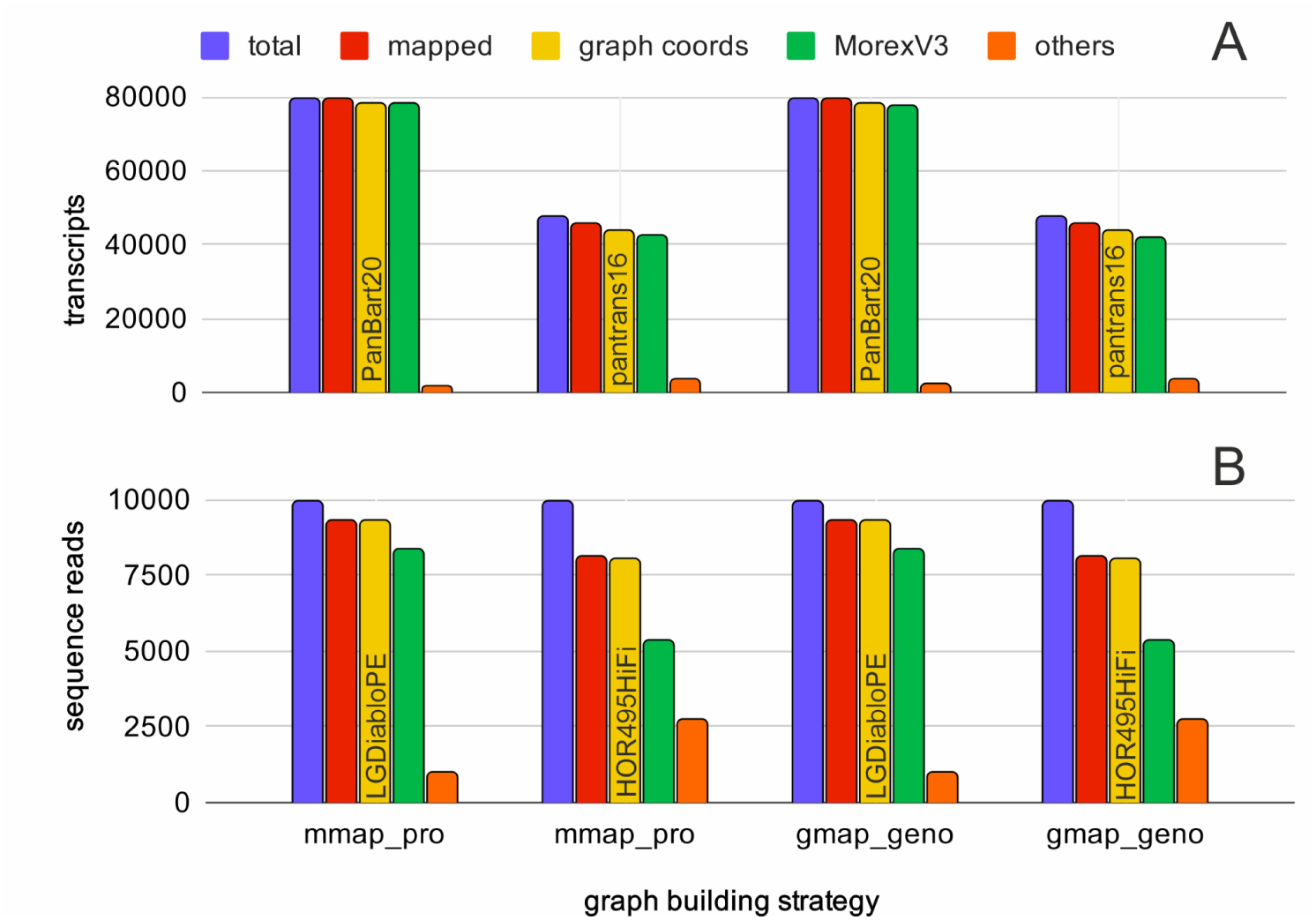
Sequences mapped to Pan20 graphs in benchmark with identity ≥ 98%, coverage ≥ 95% and range coverage ≥ 75%. A) transcripts from the *PanBaRT20* (n=79,580) and *pantranscriptome16* (n=47,588) datasets. B) genomic short and long reads datasets (*LGDiabloPE* and *HOR495HiFi*, each with 10K sequences). Mapped sequences are aligned by GMAP within MorexV3 or other genomes. Graph coord sequences are assigned graph coordinates.

Similarly, **Figure 2B** summarizes the results for short and long genomic (HiFi) sequences. In this test the mapping rate was just over 93% with short reads, and 81% for long reads, with almost all being assigned graph coordinates. Finally, we observed that 10% of *LGDiablo.PE150.10K* and 34% of *HOR495.HiFi.10K* sequences match non-reference genomes.

A different case is summarized on **Table 2**, where transcripts from three genes of cultivar Igri (HORVU.IGRI.PROJ.5HG00442610.1, HORVU.IGRI.PROJ.5HG00442620.1, HORVU.IGRI.PROJ.5HG00442640.1, see **Table S2**) are mapped to two Pan20 loci. These sequences are encoded by *CBF* loci, known to have a variable number of tandem copies across barley cultivars (Guerra et al., 2022). This example illustrates that CNV sequences, when mapped to a graph, might condense their locations on less loci than expected, particularly if extra copies can be aligned confidently to the reference genome (MorexV3 in this case). Another interesting case are sequences that are inverted in some genomes. For instance, genomic fragments containing the *HvCEN* locus in different cultivars (**Table S2**) are mapped to the same Pan20 graph location despite being inverted in cultivar Planet (**Table S3**). This example demonstrates how regions inverted in some genomes are added to the graph in the position and orientation those regions have in the landing genome, usually the reference.

**Table 2.**
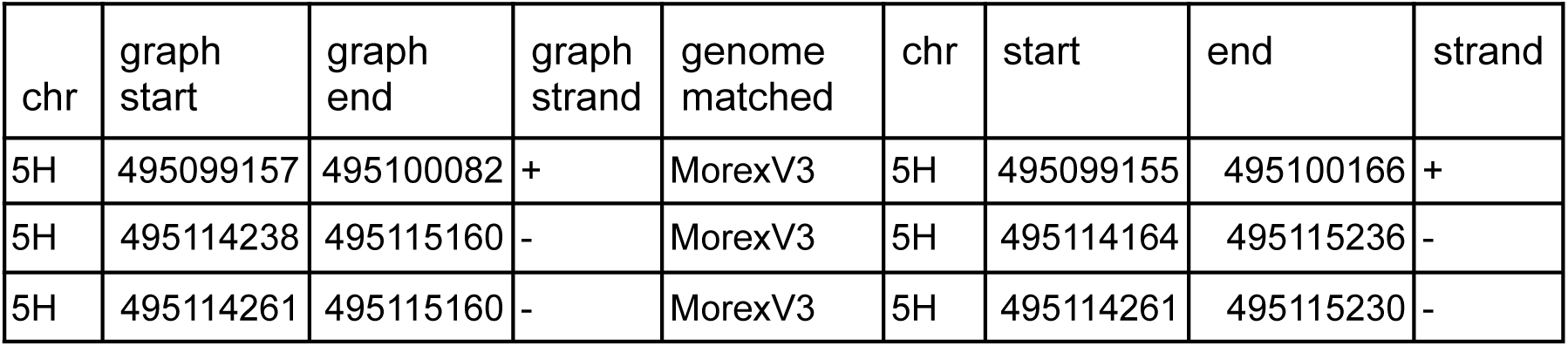
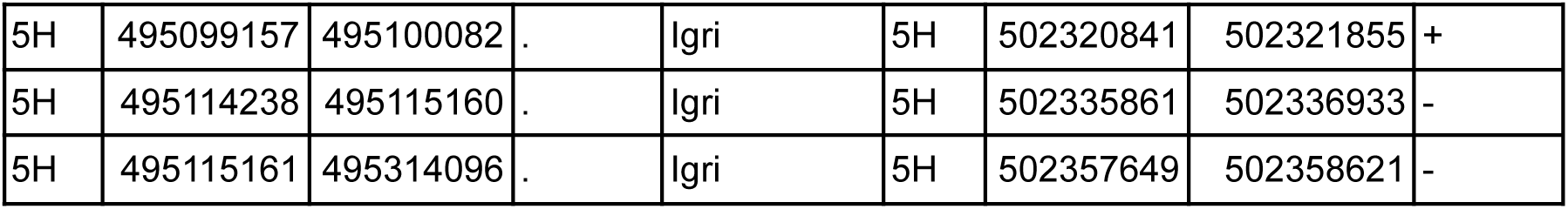
Mapping three CBF transcripts from cultivar Igri to the Pan20 graph with identity ≥ 98%, coverage ≥ 95% and range coverage ≥ 75%. Graph coordinates are shown on the left, coordinates of the first genome matched with GMAP are shown on the right. The first three rows correspond to matched locations in Pan20, with the second and third being the same locus (see graph and MorexV3 coordinates). The last three rows correspond to the same mapping with optional parameter *--single_genome Igri*, demonstrating the effect that the landing genome has on the mapping of CNV sequences.

| chr | graph start | graph end | graph strand | genome matched | chr | start | end | strand |
| --- | --- | --- | --- | --- | --- | --- | --- | --- |
| 5H | 495099157 | 495100082 | + | MorexV3 | 5H | 495099155 | 495100166 | + |
| 5H | 495114238 | 495115160 | - | MorexV3 | 5H | 495114164 | 495115236 | - |
| 5H | 495114261 | 495115160 | - | MorexV3 | 5H | 495114261 | 495115230 | - |
| 5H | 495099157 | 495100082 | . | lgri | 5H | 502320841 | 502321855 | + |
| 5H | 495114238 | 495115160 | . | lgri | 5H | 502335861 | 502336933 | - |
| 5H | 495115161 | 495314096 | . | lgri | 5H | 502357649 | 502358621 | - |

### Mapping pangenes with known coordinates

In this section we report on another benchmark where CDS nucleotide sequences from precomputed pangenes were analyzed in sequence mapping mode. In this case, the real genomic coordinates of the tested sequences was known beforehand, so it was possible to compare the landing coordinates in each graph genome to the cognate start and end positions. The results summarized in **Figure 3** and **Table S4** indicate that combining minimap and GMAP for refining range alignments (*--add_ranges both*) provides the best overall accuracy without sacrificing the number of total alignments. The default sequence identity cutoff (≥98%) retrieves the highest accurate mappings. In comparison, only the gmap_geno graph with 100% identity retrieves more alignment, but with a severe drop in accuracy (83.9% exact starts).

**Figure 3.**
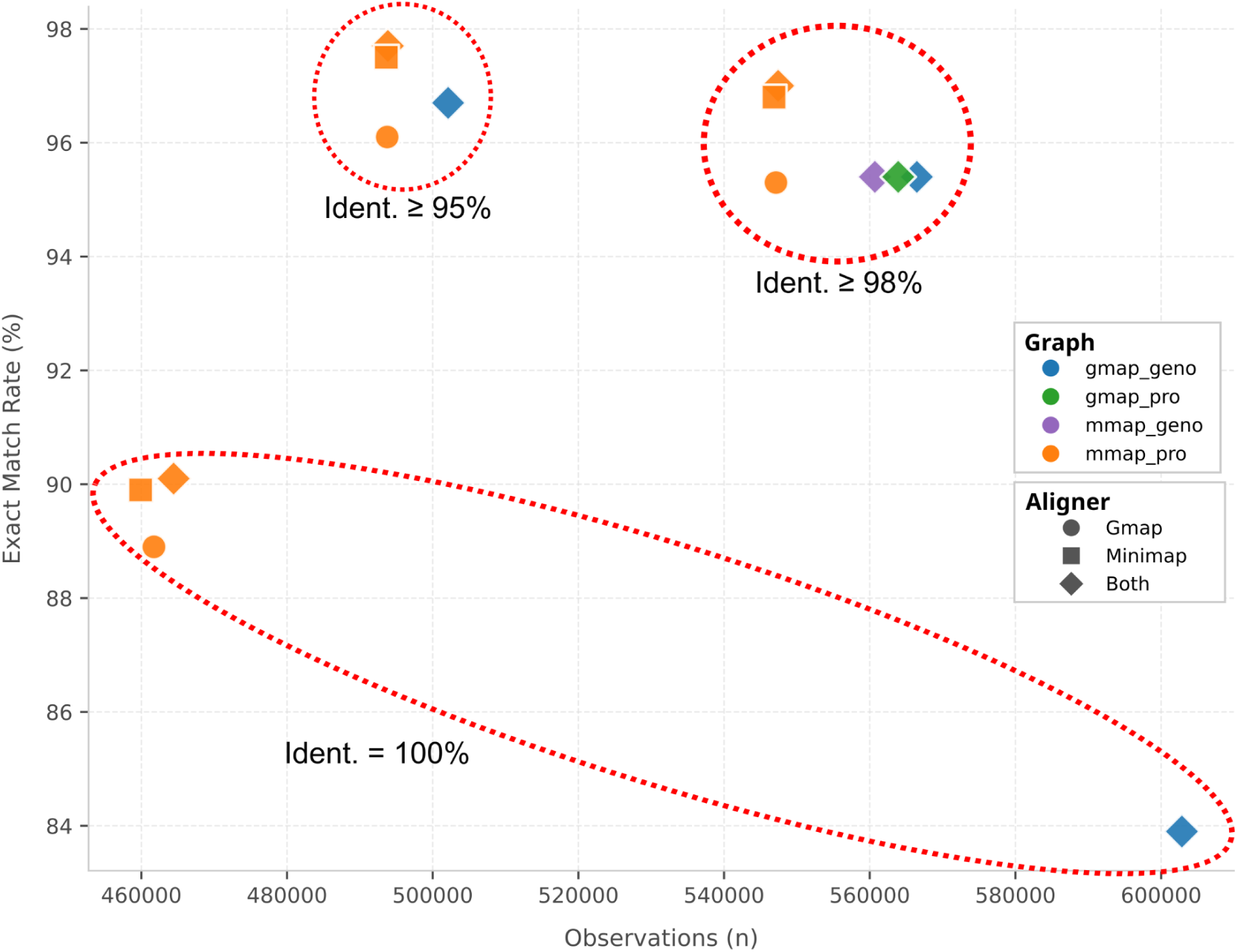
Mapping coordinate accuracy and successfully aligned pangene CDS sequences against Pan20. The Y-axis represents the percentage of exact startcoordinate match, while the X-axis shows the total number of alignments.

The data show that the *mmap_pro* graph performs better than the other, with the highest percentages of exact start (97.1%) and end (96.1%) coordinates. The gap between annotated and observed coordinates is summarized in **Figure S5**, which suggests that a few hundred base pairs are enough to collapse the curve of all genomes for 99% of the results of *mmap_proali*. In terms of successful alignments, the data indicate that the *gmap_geno* graph is slightly superior, at the cost of start/end accuracy.

### A close look at flowering pangenes

To further assess the utility of Pan20 in navigating genomic diversity we evaluated the mapping fidelity of a curated set of CDS nucleotide sequences from 28 pangenes obtained previously related to flowering and spike architecture (Contreras-Moreira et al., 2023). These genes are crucial for adaptation to diverse agroecological environments (Fernández-Calleja et al., 2021) and include allelic, CNV and PAV variation. Sequences from all genotypes included in the pangenes were mapped against alternative Pan20 graphs and the number of individual matched genomes recorded. The results in **Figure 4A** show that 22 loci can be mapped in all individual genomes by at least one Pan20 graph, while 6 are likely accessory, found only in a fraction of the genomes.

**Figure 4.**
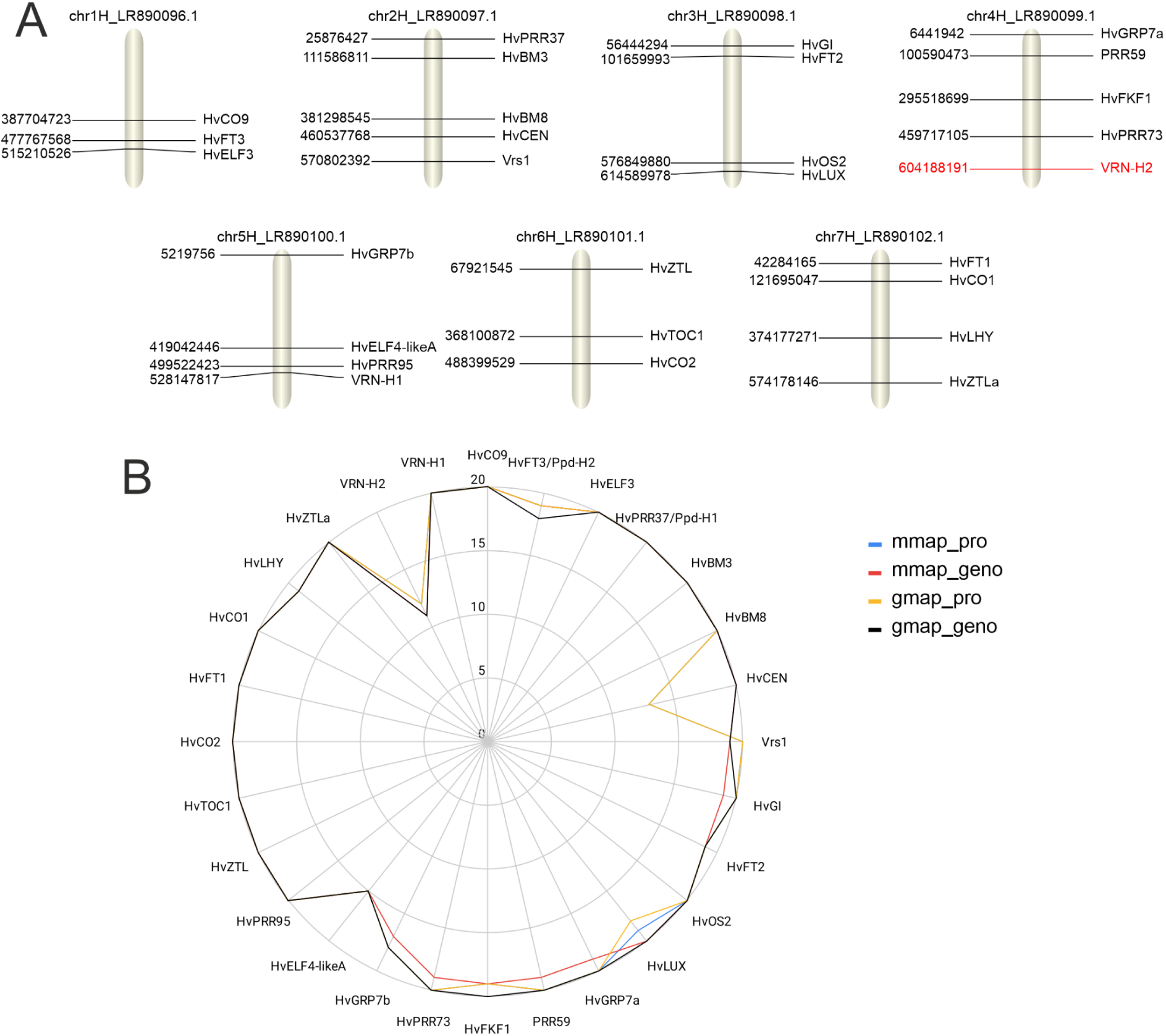
Mapping barley flowering sequences. A) Graph coordinates of 28 CDS sequences of flowering genes mapped against Pan20. The coordinates correspond to MorexV3 locations, including the estimated position of *VRN-H2*, a gene absent in the reference genome (in red). This figure was exported from the BARLEYMAP Web application at https://barleymap.eead.csic.es. B) Number of genomes in Pan20 matched after mapping CDS from 28 flowering-related genes. Four Pan20 graphs were tested: mmap_pro, mmap_geno, gmap_pro, gmap_geno.

Detailed analysis of the mapping results revealed edge cases where different lift-over and alignment algorithms used during graph construction resulted in mapping differences. As mentioned in the previous section, this analysis also exposed a trade-off between the way the graph is built and the performance when mapping genes of interest. Also, the results in **Figure 4B** show that some genes are better mapped in a particular graph, while others yield better results in a different one. An example is the *HvCEN* locus, which resides within a well-characterized inversion in chr2H shared by 7 genomes included in Pan20 (Jayakodi et al., 2020). Within the inverted genomes, graphs constructed using the AnchorWave *proali* algorithm were unable to align this locus properly. We observed the opposite behavior at the *Vrs1* locus in the OUN333 genome. Here, the sequence failed to resolve properly in graphs built with the *genoAli* algorithm, yet it was perfectly anchored when using the *proali* pipeline. Finally, analysis of the *VRN-H2* results indicate that the Pan20 graphs can assist in the analysis of locus displaying PAV and CNV events. CDS sequences encoded at the *VRN-H2* locus, which is absent from the MorexV3 genome, were successfully mapped and resolved, exposing presence-absence polymorphisms among the non-reference founder genomes and up to three copies in genotype Akashinriki.

### Graph-based imputation of GBS and low-pass data

In contrast to conventional imputation methods, which conserve empirical data and infer missing calls via population-wide linkage disequilibrium, PHG-based imputation operates through a distinct graph-centric approach. The process involves k-mer pseudo-alignment of sequencing reads against the indexed pangenome haplotype ranges. Subsequently, the optimal haplotype path, representing the most probable combination of pangenome haplotypes for the query sample, is determined by the Viterbi algorithm on the Trellis representation of the graph. The composite haplotype sequence of the optimal path is what we call the PHG-imputed sequence. In this section we map and impute GBS and low-pass data from a number of barley genotypes using Pan20 and measure its performance in two ways. In both cases we used the high quality genome assemblies of those genotypes as a control.

First, we computed the read mapping rate to measure how the choice of reference sequence affects the analysis of genotyping data. The assumption here was that using a graph as a reference would enlarge the mapping space as compared to a single reference genome such as MorexV3. Second, we wanted to assess the accuracy of barley PHG-imputation at the nucleotide level as compared to standard variant calling. The barplot on **Figure 5** (left axis) shows that k-mer pseudo-alignment against Pan20 effectively maps more GBS reads (97.71%) than simply mapping against MorexV3 (94.71%), not far from the control mapping experiment against the corresponding sample assemblies (97.78%). Regarding imputation accuracy, we took advantage of each sample assembly as the ground truth to assess variant calling fidelity. The line plots on **Figure 5** (right axis) indicate that PHG-imputed base calls have a percentage of error of 0.06%, compared to 0.32% errors of standard variant calling analysis. A closer look (**Table S5**) revealed that these estimates are mostly for SNP calls; if only indels are considered, the error rates are 3.82% and 3.09%, respectively. Also, as PHG-imputation works at the haplotype level, it can be often observed that imputation errors come in blocks (**Table S6**).

**Figure 5.**
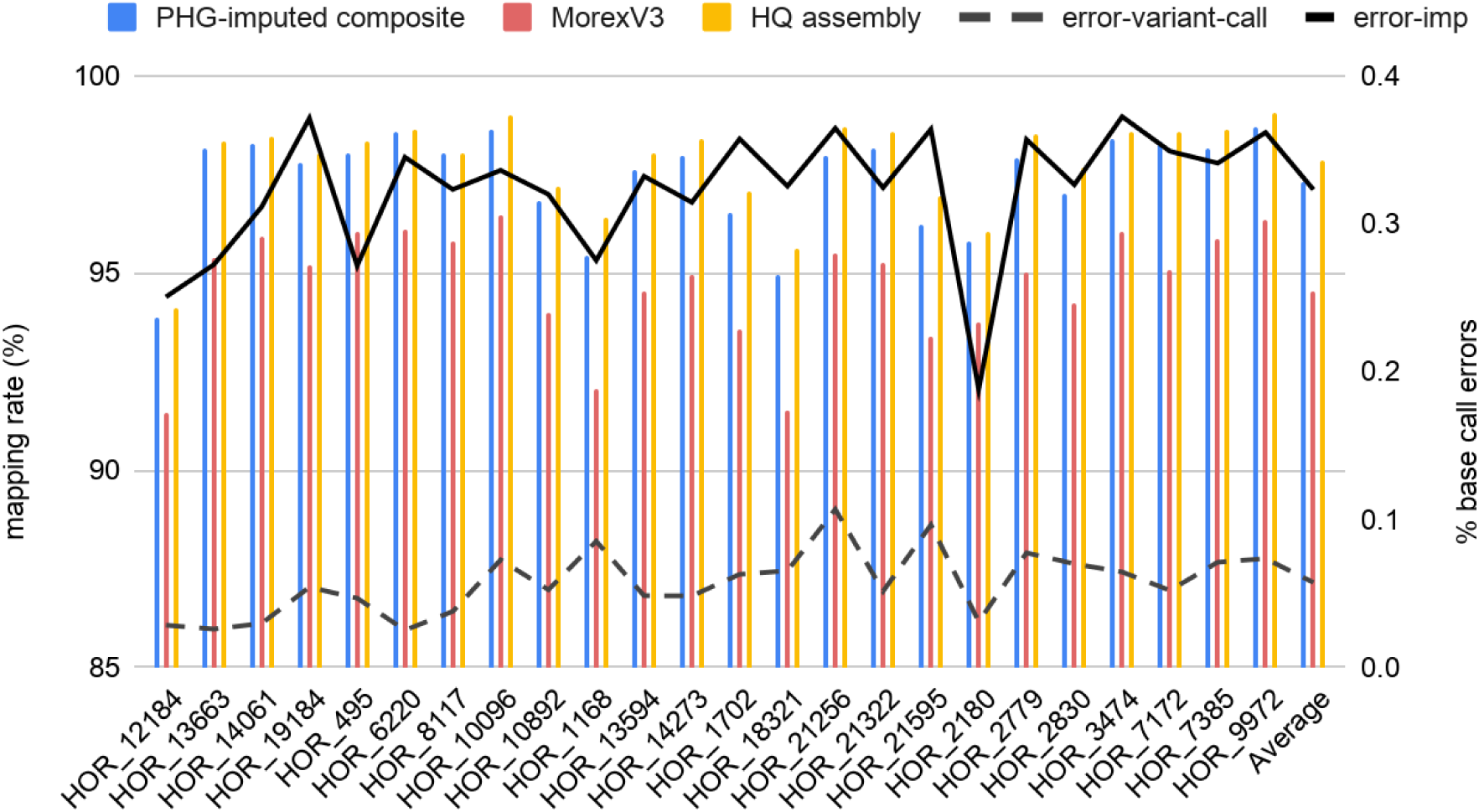
Imputation performance on Pan20 (mmap_pro) after mapping 25 GBS barley datasets from pangenome V2. Left Y-axis, columns) read mapping rate on three distinct targets (Pan20, MorexV3 and HQ assemblies). Right Y-axis) percentage of wrong base calls of PHG-imputed (dashed line) and standard variant calls (solid line, depth ≥10).

A similar test was carried out with short reads data from low-pass whole genome sequencing for 7 of the same barley genotypes. In this case the number of mapped reads is much larger. The results in **Figure S6** show that the mapping rate of k-mer pseudo-alignments is slightly higher (97.73%) than simply mapping against MorexV3 (97.45%). More importantly, the data indicate that PHG-imputed base calls have a percentage of error of 0.06%, compared to 0.36% errors of standard variant calling analysis, in line with GBS observations. Also, the error rate for indels was higher when imputing low-pass reads, with estimates of 2.61% and 1.67%, respectively (see **Table S8**).

## Discussion and conclusions

The first contribution of this paper is a protocol for building pangenome graphs for large, complex genomes such as barley. For crops this is known to be challenging due to shuffling, deletion or duplication of genes (Soltis & Soltis, 2026); in our hands, this can be done with 20 high-quality barley genomes using a conventional Linux server in a week. Of course, this was possible because we used the PHG framework, a fast and cost-effective tool for genotyping plant genomes used for instance in maize and wheat (Jordan et al., 2022). In the case of Pan20, the graphs we built contained ∼64K genomic ranges, each with 1 to 20 haplotypes, that help capture the genomic diversity of the species. When these discrete data are used to compute a haplotype-based phylogenetic tree, in particular for the Pan20 mmap_pro graph, the resulting dendrogram confirms previous results based on single-copy gene phylogenies (Chapman et al., 2026).

The lightweight data structures used by PHG support ranges and haplotypes not found in the reference genome. They are also scalable, as new genome assemblies can be added to the graph at any time. However, the reference genome still has a founder effect as i) all other genomes must be aligned to it and ii) its gene annotation defines the primary set of ranges. These limitations are partly shared with other graph architectures. For instance, a graph produced by minigraph might vary with the order genomes are added (Li et al., 2020). Other approaches such as the Variation Graph (Garrison et al., 2018) or PGGB (Garrison et al., 2024) provide intricate base-level alignment networks but quickly become computationally prohibitive in highly repetitive cereal genomes (Chapman et al., 2026).

The graphs described in this paper go beyond the PHG approach in order to optimize the process of mapping arbitrary nucleotide sequences. The *align2graph* algorithm presented here takes advantage of precomputed PHG intergenic and genic ranges and uses them to retrieve all the haplotypes that overlap matches obtained when aligning input sequences to the individual genomes making up Pan20. The alignments are computed with GMAP, an efficient aligner that can take both genomic sequences and transcripts and powers the BARLEYMAP Web application (Cantalapiedra et al., 2015). *Align2graph* is a greedy algorithm, which means sequence scans stop the moment a good match is found in a genome, saving computing time. This type of sequence search jobs, with several Pan20 graphs, can be carried out at the updated site at https://barleymap.eead.csic.es.

This hybrid mapping approach was first tested with collections of transcripts and genomic fragments. Regarding transcript isoforms, we found that 98% PanBaRT20 sequences can be confidently aligned in MorexV3, and 2% align to other genomes in Pan20. These results probably reflect the stringent filtering carried out while compiling that dataset (Guo et al., 2025). These figures shift to 93% and 7% when isoforms from the dataset pantranscriptome16 are mapped. In order to do a similar experiment with genomic sequences we aligned short and long reads. We found that 10-34% of sequences could not be mapped with default sequence identity and coverage against MorexV3. Overall, these experiments measure how much mapping space is gained when a pangenome is used as a reference instead of a single genome.

An alternative mapping experiment was carried out with CDS sequences extracted from precomputed pangenes from the same set of barley genomes (Contreras-Moreira et al., 2023). After trial and error and benchmarking, we found two main ways of building a graph that work on barley. While the Pan20 gmap_geno graph was able to map most CDS sequences, Pan20 mmap_pro was overall better, balancing the number of successful alignments and the accuracy of their coordinates. Moreover, the experiment with flowering genes revealed that mapping accuracy often depends on the underlying graph topology. In our hands no single lift-over mode could resolve all structural variations present, underscoring the need to test multiple graph-building strategies. The CBF example also revealed that in loci with CNV the order of genome scanning might have an effect on the final map produced. This is a limitation of the greedy approach tested here, but one that might be possible to avoid by changing the scanning order of genomes (supported in Docker container uniquely). If only the MorexV3 reference was to be scanned, this limitation would apply to all CNV cases where the number of copies in MorexV3 is smaller than in the genome of interest.

In order to evaluate the performance of the haplotype analysis mode we tested the k-mer mapping utilities builtin in PHG with GBS and low-pass FASTQ files. In particular we used data from genotypes not used in the construction of Pan20 graphs, and compared the obtained haplotypes with base calls obtained from standard variant calling. Because PHG relies on matching contiguous ancestral haplotype blocks rather than population-wide linkage disequilibrium, the expectation was that it would be better identifying alleles present in the graph, particularly those absent from the MorexV3 reference (Bradbury et al., 2022). This is in contrast to LD-based imputation tools like Beagle (Browning et al., 2018), that have been reported to struggle while imputing rare alleles in sparse agricultural datasets (Long et al., 2022). The GBS experiments (approximately 1M reads per sample) demonstrated that the mapping rate of Pan20 is better than the control MorexV3 mapping, as expected. Moreover, the results show that graph base calls had consistently a lower error rate than simply mapping to the reference genome. While the mapping rate improvement was modest in the low-pass experiment (hundred million reads per sample), again we were able to observe a smaller base call error when using the Pan20 graph, particularly for SNP calls. These results would agree with previous reports on other crops that suggested a superior imputation accuracy of PHG over standard linkage disequilibrium-based tools, particularly when utilizing sparse, low-coverage datasets such as GBS (Franco et al., 2020), low-pass Whole Genome Sequencing data (Jensen et al., 2020) or exome capture data (Wang et al., 2023). However, we anticipate two cases where the imputation performance would drop. The first would be sequencing samples that fail to cover equally all genomic regions, which would compromise the k-mer based algorithm, leading to spurious haplotype assignments and ultimately erroneous variant calls. A second edge case are samples that carry truly novel haplotypes not yet included in the graph. Because the Viterbi algorithm cannot construct novel paths for unrepresented germplasm, original sequences would be erroneously substituted with the most genetically similar haplotypes present in the graph. This potential data loss is the trade-off for significantly reduced computational overhead. However, as new genome assemblies become available and are added to the graph, the probability of accurately calling novel alleles will improve.

In summary, we present a graph-based framework that combines the ability of GMAP to produce compex, intron-aware alignments in complete genomes with pangenome ranges precomputed with PHG, significantly reducing the time needed by users to scan large numbers of genomes, twenty in the case of Pan20. We expect this novel strategy will prove useful in other pangenomes. Furthermore, in our tests for the tasks of mapping and imputation, it proved to be accurate and scalable. Accurate, for it can efficiently map genomic and transcriptomic sequences with correct and consistent start and end coordinates, even for loci not found in the reference genome. Scalable, as adding new high quality genomes to the graph has a linear computational cost. As barley has a complex, large genome, we expect the approach described in this paper will potentially benefit researchers of other crops, helping them explore genetic diversity beyond a single reference.

## Acknowledgements and funding

We thank the authors of PHGv2 for their feedback, patience and support on using their software and Rubén Sancho for advice on phylogenetic analyses. This work was supported by AEI/10.13039/501100011033/FEDER/UE [PID2022-142116OB-I00 and predoctoral contract PREP2022_EEAD_52 to JSA], Horizon 2020 PRIMA [PCI2019-103526] and SusCrop ERA-NET Recobar [771134], Government of Aragon [A08_23R] and CSIC [FAS2022_052, INFRA24018].

## Author contributions

Funding was acquired by AMC, EI and BCM. Methodology was designed by BCM, AMC, HA and JSA. Web programming was performed by CJR. Data was analysed by JSA, HA, AMC, EI and BCM. Research was supervised by AMC, EI and BCM. Summarized data and graphics were created by JSA and BCM. The original draft was prepared by JSA, AMC and BCM. All authors revised and approved the final manuscript.

