## Supplementary figures for "A pangenome-graph approach for mapping and imputing barley sequences"

Estación Experimental de Aula Dei-CSIC, Av. Montañana 1.005, 50059 Zaragoza, Spain

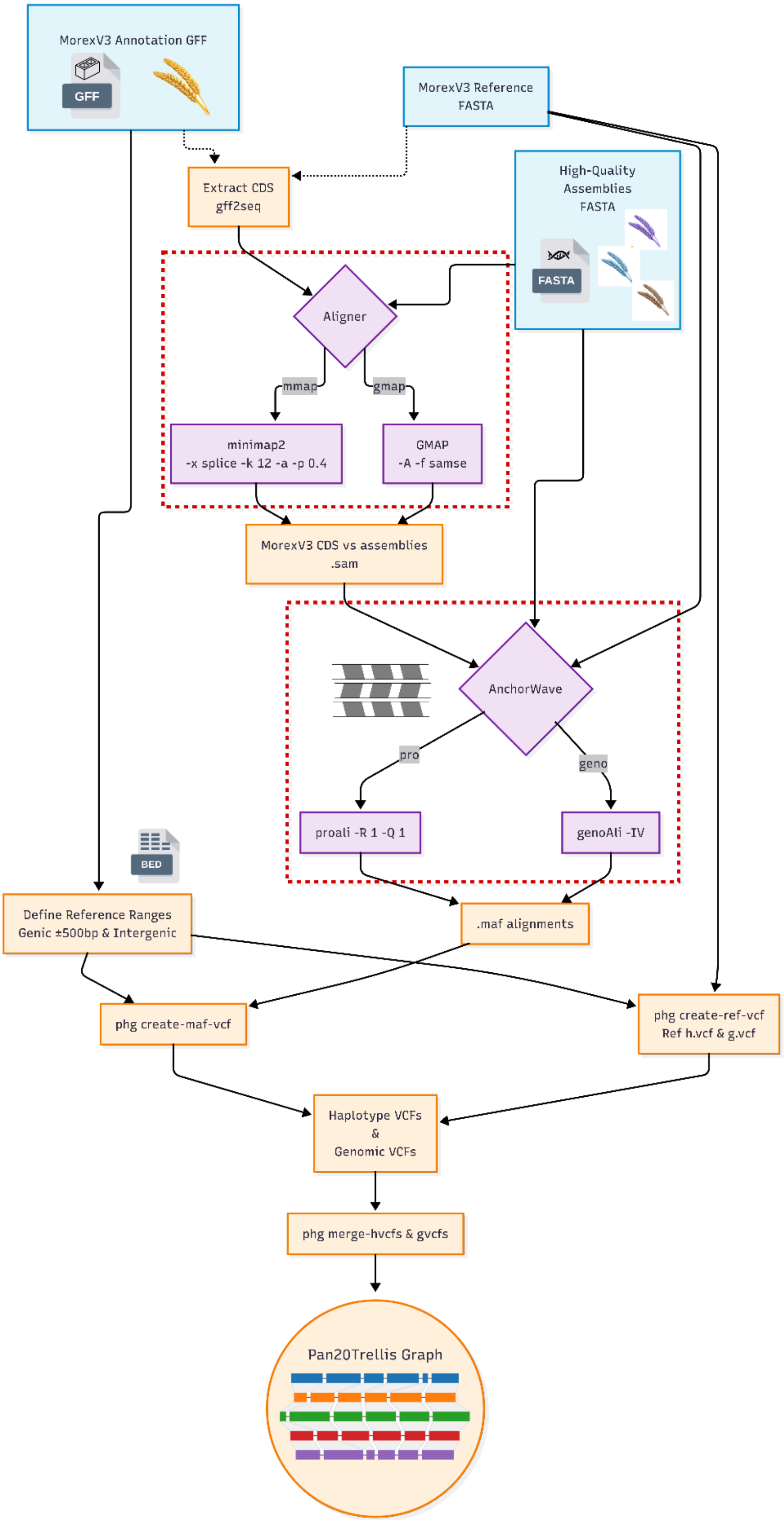


**Figure S1**. Protocol for building a PHG-based pangenome graph. 1) The graph is seeded with the assembly (FASTA) and the gene annotation (GFF) of the reference genome, MorexV3 in this work. Reference gene models contribute the first set of genomic ranges to the graph, which are consecutive genic and intergenic regions. 2) Other high-quality assemblies are added one-by-one, aligned to MorexV3 with the AnchorWave algorithm, adding new ranges not previously considered to the graph (see Methods). 3) The resulting graph maps ranges across genomes, supporting also presence-absence variation, and can be transversed to impute haplotypes.

**Figure S2**. Barleygraph align2graph flowchart.

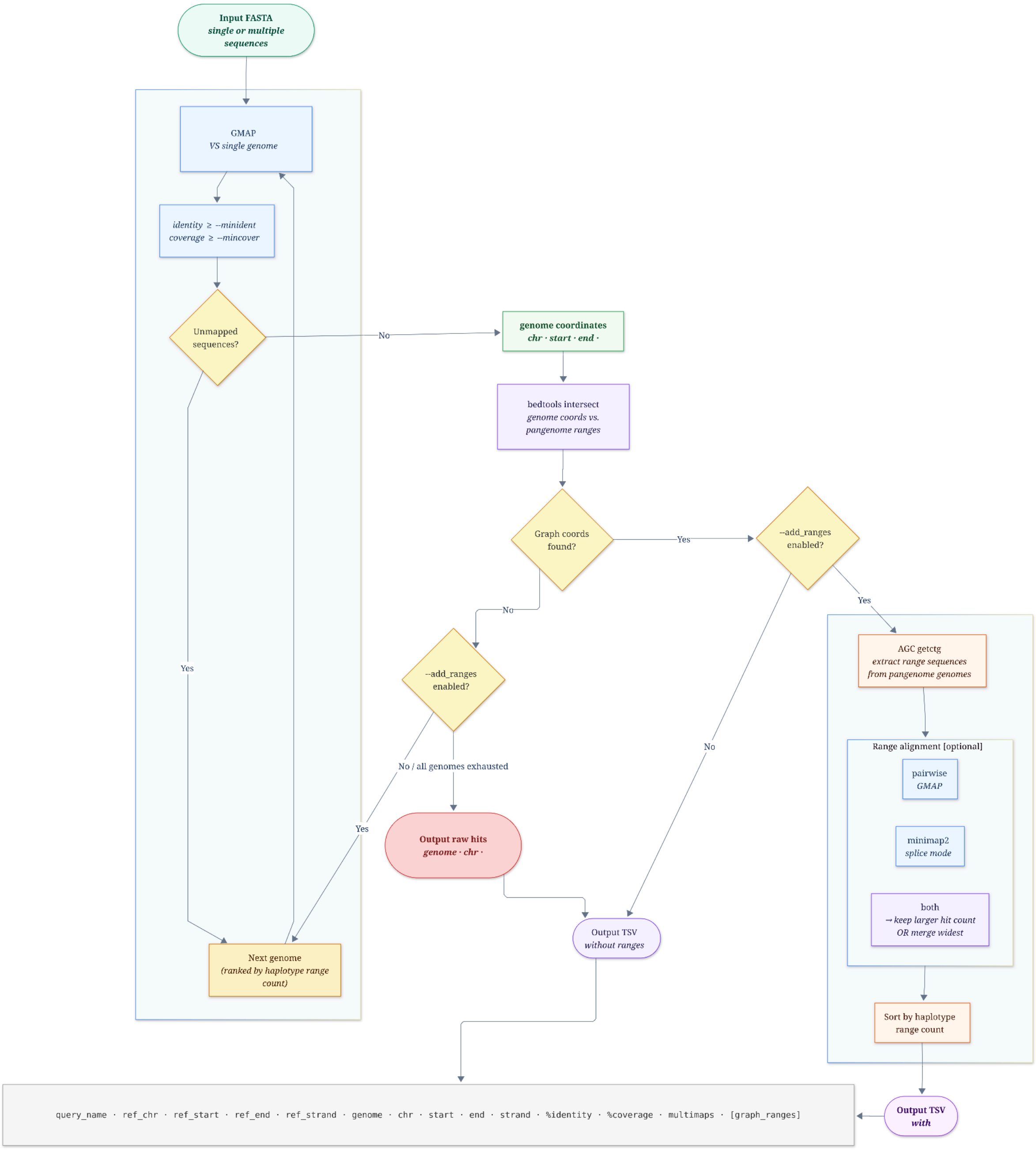

**Figure S3**. Updated BARLEYMAP Web application. A) Two types of input can be queried: identifiers/query IDs and nucleotide sequences. B) The “Graph” mode/*align2graph* (green box on the right) scans input FASTA sequences hierarchically with GMAP, starting with MorexV3 and moving to the next genome in the graph until a match satisfying the coverage and identity cut-offs, and overlapping graph ranges, is retrieved. If there are several overlapping ranges (two in Du_Li_Huang in the example), the range with most genomes is selected. Finally, the coordinates of the corresponding graph ranges are refined by aligning the input sequences against them with GMAP and minimap2.

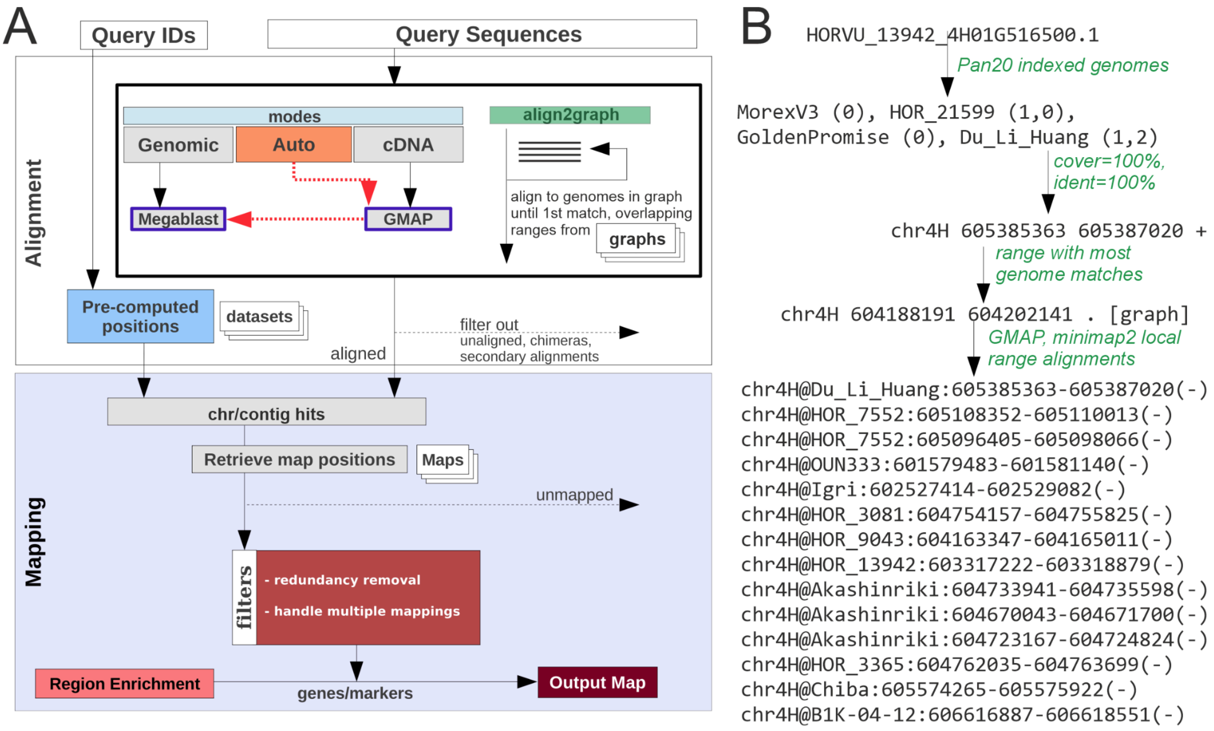

**Figure S4**. Example of 'Graph' results in BARLEYMAP (https://barleymap.eead.csic.es). Top) Diagram showing the location of input sequences in the graph. Middle) global graph coordinates of input sequence (VRN2 in the example). If sequence is absent in the MorexV3 reference the strands is shown as a dot ".". The "other alignments" columns lists the estimated genomic coordinates of the input sequence in all graph genomes where a good match was found; there might be several hits on the same genome. Bottom) Table with markers in BARLEYMAP in the vicinity of the graph coordinates of mapped input sequences, including MorexV3 genes. In cases where precomputed pangenes can be inspected, they appear as clickable markers.

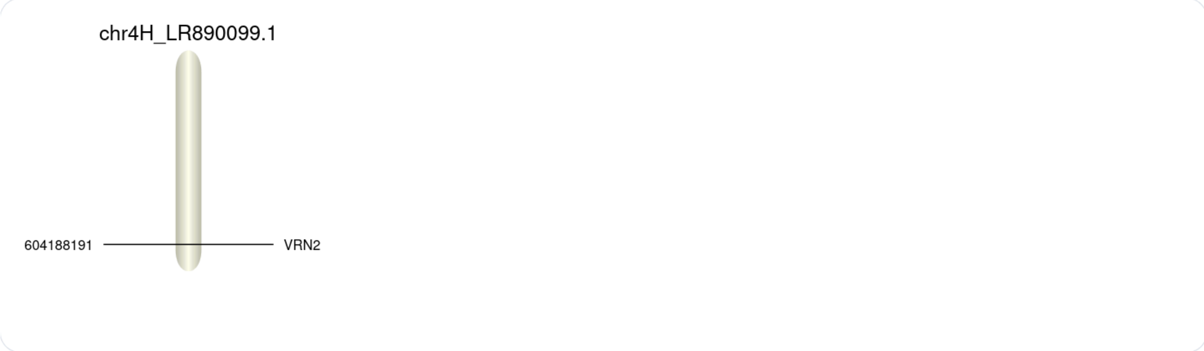


| Marker | chr | start | end | strand | other alignments |
|---|---|---|---|---|---|
| **VRN2** | **chr4H_LR890099.1** | **604188191** | **604202141** | **.** | **chr4H@HOR_9043:604163347-604165011(-)<br>chr4H@OUN333:601579483-601581140(-)<br>chr4H@Igri:602527414-602529082(-)<br>chr4H@Du_Li_Huang:605385363-605387020(-)<br>chr4H@B1K-04-12:606616887-606618551(-)<br>chr4H@HOR_13942:603317222-603318879(-)<br>chr4H@HOR_7552:605108352-605110013(-)<br>chr4H@HOR_7552:605096405-605098066(-)<br>chr4H@HOR_3365:604762035-604763699(-)<br>chr4H@HOR_10350:608218048-608219712(-)<br>chr4H@Akashinriki:604733941-604735598(-)<br>chr4H@Akashinriki:604670043-604671700(-)<br>chr4H@Akashinriki:604723167-604724824(-)<br>chr4H@Chiba:605574265-605575922(-)<br>chr4H@HOR_3081:604754157-604755825(-)** |

| Marker | chr | start | end | strand | other alignments | Description | GeneOntologies |
|---|---|---|---|---|---|---|---|
| BART1_0-u31967 | chr4H_LR890099.1 | 604184122 | 604188347 | - | No | | |
| **VRN2** | **chr4H_LR890099.1** | **604188191** | **604202141** | **.** | **No** | | |
| MLOC_22857 | chr4H_LR890099.1 | 604194335 | 604195591 | - | No | | |
| HORVU4Hr1G088500.2 | chr4H_LR890099.1 | 604194752 | 604195123 | - | No | | |
| BaRT2v18chr4HG219170.1 | chr4H_LR890099.1 | 604196710 | 604197817 | - | No | | |
| PanBaRT20_chr4HG43155.1 | chr4H_LR890099.1 | 604196815 | 604197981 | - | No | | |
| PanBaRT20_chr4HG43156.1 | chr4H_LR890099.1 | 604196949 | 604198098 | + | No | | |
| BaRT2v18chr4HG219180.1 | chr4H_LR890099.1 | 604197128 | 604197809 | + | No | | |
| HORVU.MOREX.r3.4HG0416230 | chr4H_LR890099.1 | 604197212 | 604197799 | + | No | Virulence factor-related M protein | GO:0005515 GO:0005576 GO:0005618 GO:0006909 GO:0009405 GO:0016020 |
| HORVU4Hr1G088510.1 | chr4H_LR890099.1 | 604199975 | 604200154 | - | No | | |
| BaRT2v18chr4HG219190.1 | chr4H_LR890099.1 | 604202030 | 604217298 | + | No | | |
| PanBaRT20_chr4HG43157.1 | chr4H_LR890099.1 | 604202072 | 604217166 | + | No | | |
| BART1_0-u31956 | chr4H_LR890099.1 | 604202089 | 604217349 | + | No | | |

**Figure S5**. Accuracy of start and end coordinates assigned to CDS sequences from barley pangenes mapped to Pan20 (mmap_pro) with default coverage and identity (95% and 98%) and parameter *--add_ranges both*. The Y-axis shows the percentage of mapped sequences displaying the difference between known and mapped coordinates shown in the X-axis. The total number of mappings for each genome in the graph is shown in the legend.  Sequences from the HOR_10350 genotype were excluded due to a genome version mismatch between the pangenes set and Pan20.

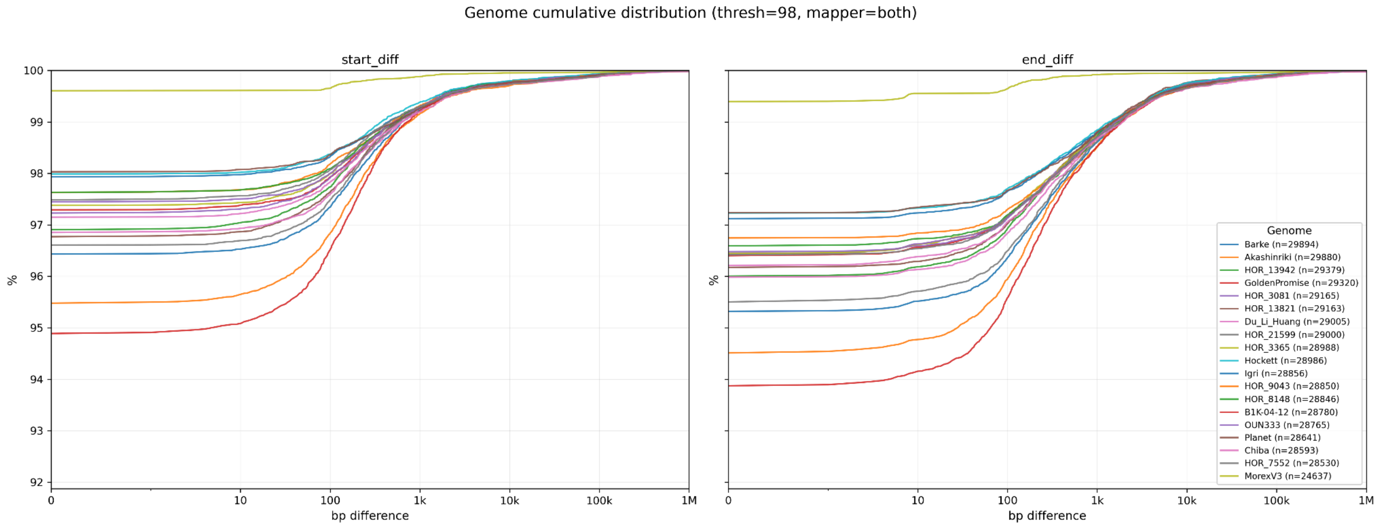

**Figure S6**. Imputation performance on Pan20 (mmap_pro) after mapping 7 low-pass genome sequencing datasets. Left Y-axis, columns) read mapping rate on three distinct targets (Pan20, MorexV3 and HQ assemblies). Right Y-axis) percentage of wrong base calls of PHG-imputed (dashed line) and standard variant calls (solid line, depth ≥5).

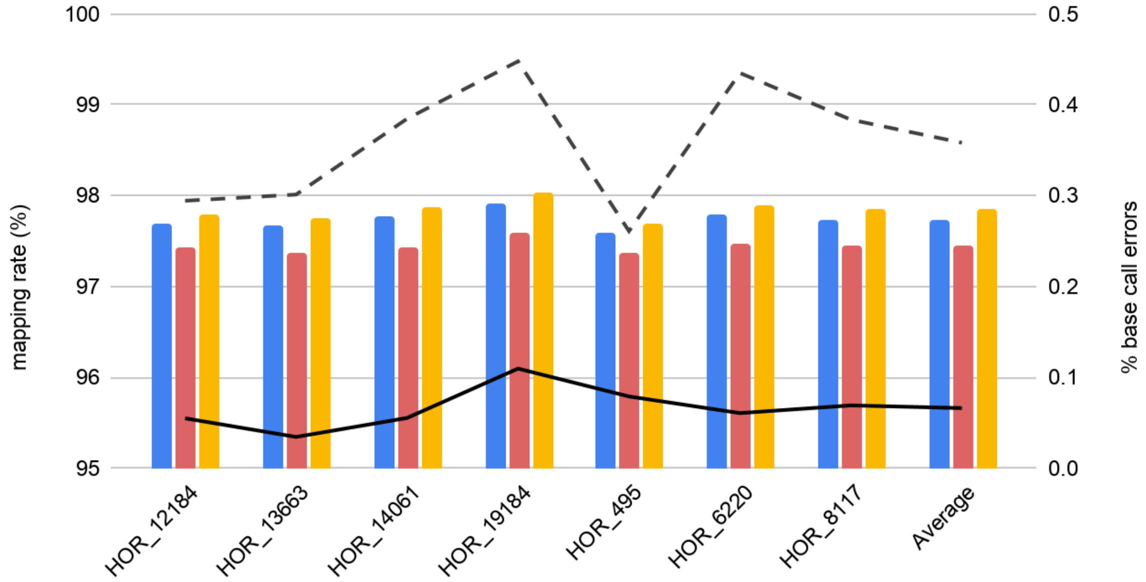
